# Changes in large-scale neural networks under stress are linked to affective reactivity to stress in real life

**DOI:** 10.1101/2023.03.28.534537

**Authors:** Rayyan Tutunji, Martin Krentz, Nikos Kogias, Lycia de Voogd, Florian Krause, Eliana Vassena, Erno J. Hermans

## Abstract

Controlled laboratory stress induction procedures are very effective in inducing physiological and subjective stress. However, whether such stress responses are representative for stress reactivity in real life is not clear. Using a combined within-subject functional MRI laboratory stress and ecological momentary assessment stress paradigm, we investigated dynamic shifts in large-scale neural network configurations under stress and how these relate to affective reactivity to stress in real life. Laboratory stress induction resulted in significantly increased cortisol levels, and shifts in task-driven neural activity including increased salience network (SN) activation in an oddball task and decreased default mode network activity in a memory retrieval task. Crucially, individuals showing increased SN reactivity specifically in the early phase of the acute stress response also expressed increased affective reactivity in real life. Our findings provide (correlational) evidence that real-life affective stress reactivity is driven primarily by vigilant attentional reorienting mechanisms associated with SN.

## Introduction

Acute stress triggers a cascade of time-dependent processes that result in dynamic shifts of large-scale brain network configurations ^1,2^. These processes are driven by distinct actions of stress-sensitive hormones and neuromodulators in the acute phase and in the recovery phase of the stress response ^3,4^. While these dynamics have been studied extensively in experimental models in both animals and humans, it is unknown how interindividual differences in stress-induced perturbations of large-scale networks are reflected in interindividual differences in reactivity to acute stressors in real life. This variability is critical for understanding the role of networks/stress responses in healthy adaptation to stress and psychopathology, especially given the involvement of alterations in large-scale networks in both stress and psychopathology ^5–8^.

Research has implicated three core large-scale brain networks - the salience (SN), executive control (ECN), and default mode networks (DMN) - in both reactivity to, and recovery from a stressor through the effects of stress-related hormones and neurotransmitters ^1,6^. The early reactivity phase of the stress response is thought to be driven by actions of catecholamines and corticosteroids ^4,9^. In this phase, activation of the locus coeruleus results in tonically elevated release of norepinephrine ^10^. Increased LC activity and norepinephrine have also been associated with overall increased salience network activity ^3,11,12^. This coincides with specific environmental demands during acute stress, with an increased need for threat vigilance ^13^. At the same time, studies have also shown ECN and DMN suppression, associated with impaired working memory and memory retrieval under acute stress, functions supported by these two networks, respectively ^14–16^.

Interestingly, these processes may be reversed through slow, gene transcription-dependent corticosteroid effects in the later recovery stage of the stress response, starting 1-2 hours following stress ^4^. Following the administration of hydrocortisone, within a time window in which genomic actions can be expected, a decrease in SN-related regions is seen ^17,18^. Delayed effects of hydrocortisone in a time window that is consistent with genomic mechanisms have also been linked to upregulation of regions associated with the ECN, as well as with improved working memory performance ^19^. Additionally, delayed effects of stress hormones have been shown to result in improved DMN linked memory retrieval ^20^. Thus, growing evidence points to a reversal of overall changes in network balance under stress, with a decrease in SN, and increase in DMN and ECN related regions driven by later, genomic effects of corticosteroids. Arguably, this reversal in the late stage of the stress response serves an adaptive function by promoting higher-order cognitive functions required for a return to homeostasis and an optimal preparation for future stressors. While many studies have shown the shifts within these large-scale networks over the course of either the early or late stress response, the temporal dynamics have thus far been only inferred. Within-subject, time dependent shifts in these large-scale networks over the full duration of the stress response – including both early nongenomic and late genomic effects – have not been thoroughly investigated.

While laboratory findings are important in establishing a mechanistic understanding of the neural stress response, a critical question is how individual dynamics of neural stress reactivity relate to healthy and functional responses to stress in real life. Real-life stress is often studied using approaches such as Ecological Momentary Assessments [EMA, also known as experience sampling methods or ESM^21^]. These methods leverage repeated assessments in day-to-day lives of participants to derive measures of stress reactivity and sensitivity ^22–24^. While such studies have created the opportunity to quantify stress reactivity in daily life, linking these observations to the lab has been a more difficult endeavor. Current attempts to derive daily-life stress measures are limited by study designs that are unable to disentangle measures of exposure to stressors from measures of the consequences of said exposure. One measure of reactivity that can be adapted from resilience research to EMA studies is the residualization-based stressor reactivity measure ^25^. In resilience research, this measure is derived by regressing the change in mental health onto a measure of exposure to life stressors. The residuals of this regression then indicate changes in mental health that are not explained by the amount of stress exposure ^26–28^. Here, we adapted this measure for EMA research to quantify individual affective reactivity to acute stressors in real life at a time scale that is comparable to laboratory studies on acute stress ^29^. As an outcome measure, we used changes in positive affect, which have directly been linked to resilience ^30^.

In this study, we used a within-subject cross-sectional design (n=83) to investigate the effects of an established psychosocial laboratory stressor on the dynamics of large-scale networks under stress and task demand (see Figure 1). We investigated both the initial reactivity (i.e., early phase) and recovery phases (i.e., late phase) of this response in a novel functional MRI paradigm combining three tasks (facial oddball, numeric 2-back, and associative memory retrieval) that have each been shown to preferentially recruit one of the three core large-scale networks of interest (SN, ECN, and DMN, respectively). We then compared temporal dynamics under stress to a control scan using a matched, non-stressful procedure to determine within-person changes in large-scale network balance across the phases of the stress response. Next, we investigated the relationship between this balance and real-life affective reactivity to stress. In addition to the two scan sessions, all participants underwent two weeks of EMA, one during a high-stakes exam week, and one during a control week. We used the residualization-based approach explained above to compute stressor reactivity scores. We expected to see increased SN, and decreased ECN and DMN activity during the early phase of the stress response, with a reversal of this balance in the later recovery phase (i.e., decreased SN, and increased ECN and DMN). This hypothesis is based on a previously published working model^1^. We also expected greater SN activation and ECN and DMN suppression to be associated with increased real-life stress reactivity, with the opposite effects in the recovery phase of the neural response to stress (i.e., smaller decreases or recovery in SN and increases in ECN and DMN).

**Figure 1.**
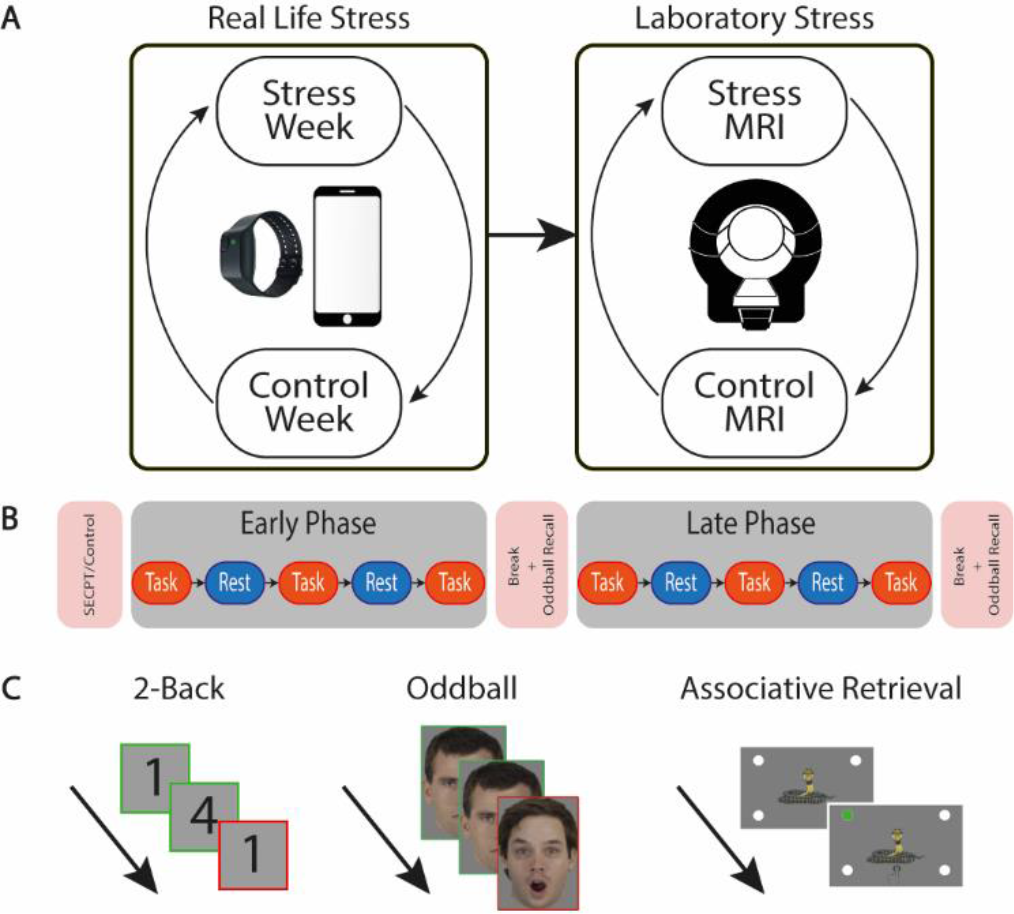
Study Design. A) Stress and control weeks were counterbalanced for order, followed by stress and control MRI scans that were also counterbalanced between subjects. B) Typical scan day for a participant. C) Three tasks performed during MRI task blocks including a standard 2- Back, facial oddball, and associative retrieval task.

## Results

### Deriving a residual-based measure of affective reactivity to stress exposure

In order to establish the validity of our real-life stress paradigm, we first verified using EMA data whether exam weeks were more subjectively stressful than control weeks without exams. During these weeks, students received repeated assessments (i.e., beeps) probing perceived stress exposure and affect. We used a self-report measure of perceived stress exposure combining three types of stress exposure: Event-related (probing the most significant event since the last beep), activity-related (probing the most significant current event), and social stress. Positive affect was measured using a four-item questionnaire adapted from previous EMA studies (see Tutunji et al. 2023 for details ^31^). Participants showed a significant increase in the number of beeps during the exam week in which perceived stress exposure was above their overall average (Odds Ratio=-1.67, SE=0.14, t-stat=6.033, p<0.001, Figure 2A) in addition to lower overall positive affect (β=-0.77, SE=0.11, t-stat=-6.80, p<0.001, Figure 2B) compared to the control week.

**Figure 2.**
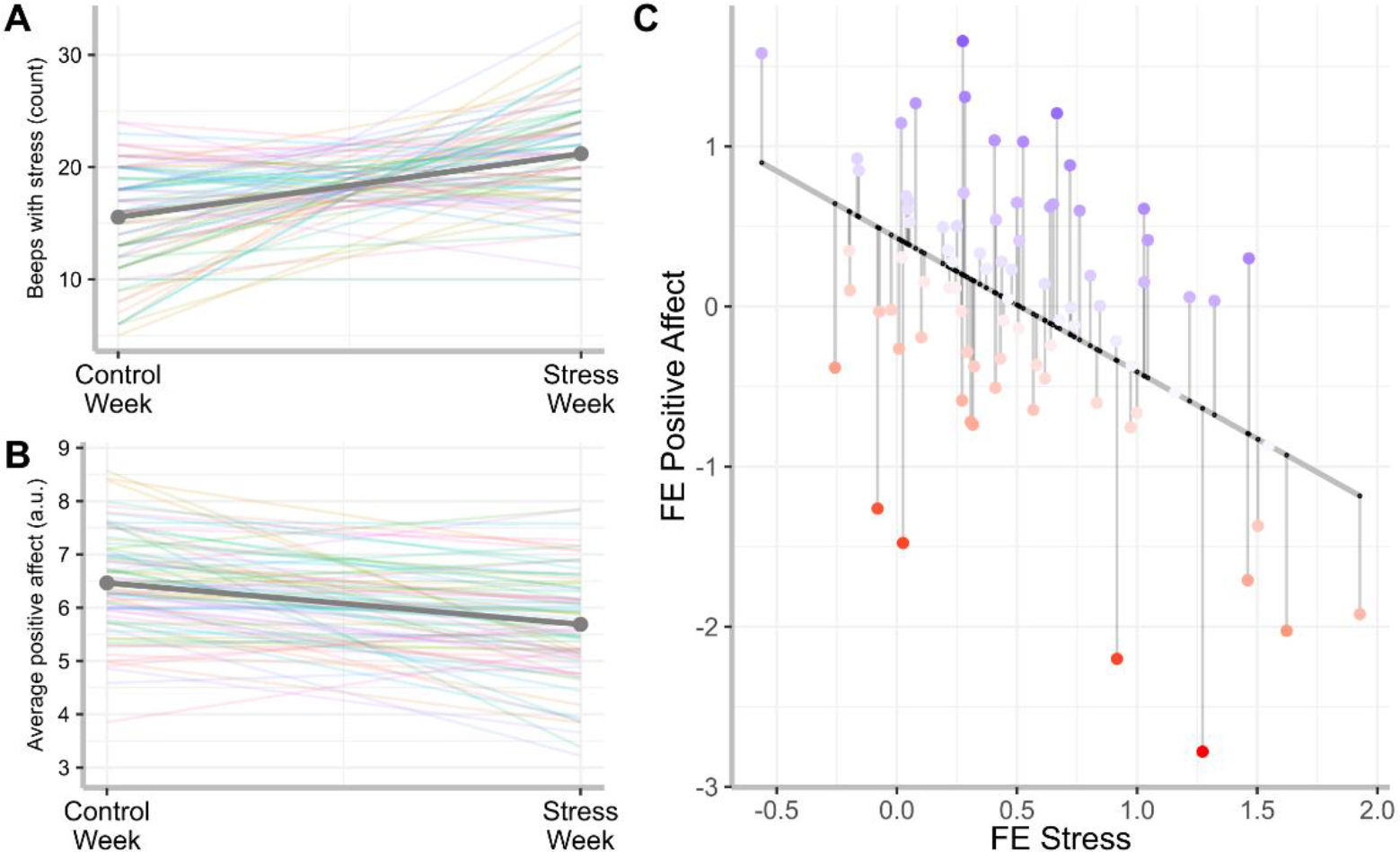
**Residualization-based affective reactivity score calculation**. The averaged effects of examination weeks on (A) subjective stress levels and (B) positive affect, with individual colored lines shown for the modeled random subject-level effects. (C) These random effects (RE) were extracted from each model and then regressed onto each other. Positive affect changes were then residualized with respect to the exposure measure and inverted, resulting in affective reactivity scores. Thus, blue dots above the line indicate lower affective reactivity to stress exposure, while red dots under the regression lines indicate higher affective reactivity scores.

To derive subject-level estimates of the effects of exam weeks on mood and perceived stress exposure, random effects were extracted from the mixed effects models. We found a significant negative association between increased stress exposure and reduced positive affect in the exam week as compared to the control week (β=-0.84, SE=0.17, t-stat=-5.09, p<0.001, Figure 2C). By extracting the residuals from this analysis, a measure can be derived of the deviation from the expected change in positive affect relative to the amount of reported stress exposure. That is, residuals above the line indicate a lower impact of stress exposure on affect, and vice versa for those below the line. The inverse of these residuals can thus be used as a residual-based affective reactivity score to estimate the within-subject reactivity controlling for potential differences in individual levels of perceived exposure to stress (Figure 2C). As a final step, we investigated whether interindividual differences in stress reactivity were related to personality or trait characteristics that are linked to psychopathology. Therefore, we correlated this measure with neuroticism scores from the NEO-FFI. There was a trend-level association between neuroticism and the residual-based score (r(72)=0.21, p=0.054).

### Cortisol responses and autonomic responses to the SECPT

We next investigated whether our laboratory stressor was effective in inducing changes in cortisol and autonomic arousal. Results of the mixed model showed a significant stress (stress, control) by time (1-5 samples at T1=-3, T2=14, T3=42, T4=87, and T5=160 min relative to stress onset) effect on salivary cortisol levels (log transformed estimate, β=1.00, SE=0.006, t-stat=2.44, p=0.015, Figure 3A). There was an additional significant effect of sex (β=-0.16, SE=0.08, t-stat=-2.06, p=0.040), with the difference being driven by attenuated cortisol responsiveness in hormonal contraceptive users specifically (β=0.26, SE=0.10, t-stat=2.69, p=0.007). Follow-up tests showed significantly lower cortisol in the stress session relative to the control session immediately following the stress induction procedure, after Tukey correction (T=9 minutes, β=0.173, SE=0.084, p=0.039), but expectedly higher cortisol at the third and fourth samples (T=39 minutes, β =-0.296, SE=0.84, p<0.001, and T=85 minutes, β=-0.165, SE=, p=0.0497). No significant differences in cortisol were observed in the last sample (T=160 minutes, SE=-0.101, t-stat=0.084, p=0.229). We used the difference in the area under the curve with respect to increase (AUCi) in the early phase as a base measure of cortisol stress reactivity in later models. To this end, a mixed effects model showed that AUCi of salivary cortisol was significantly higher in the stress session compared to the control session, when controlling for scan order effects (Mdiff=103.86, t=3.975 p<0.001).

**Figure 3.**
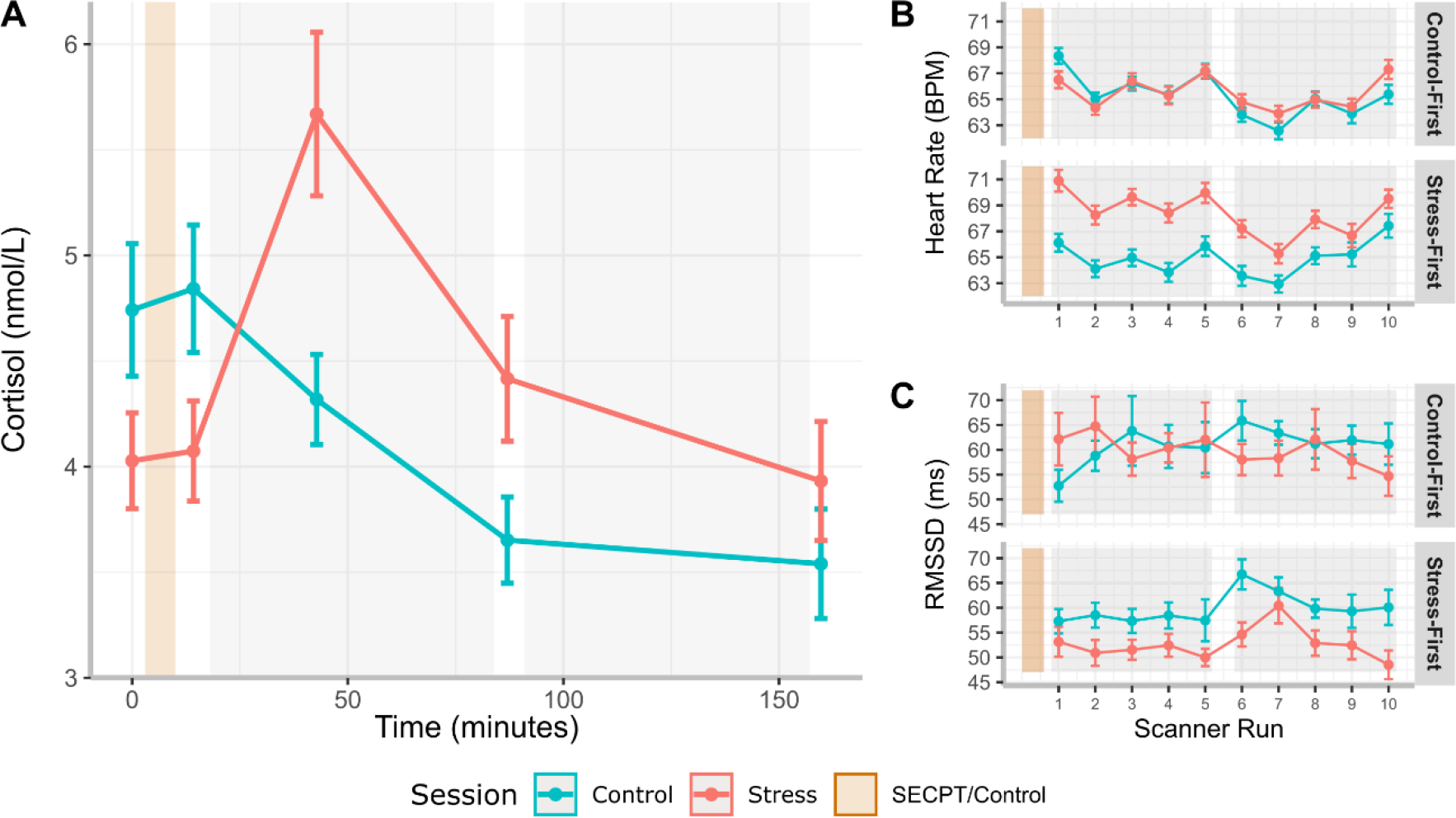
Physiological changes in response to SECPT. A) Salivary cortisol response showing increased cortisol in response to the SECPT compared to the control procedure. B) Average heart rate (beats per minute) and C) average heart rate variability (RMSSD, ms) per scanner run separated by whether participants had the control or stress scan first. Participants who had the stress session first exhibited increased heart rate and decreased heart rate variability (RMSSD) during the stress session. Error bars = SEM.

We furthermore examined autonomic reactivity to stress by looking at average heart rate and heart rate variability during the scanner runs (starting at 18, 30, 41, 53, 60 minutes post stress in the early phase, and 98, 111, 117, 128, and 135 minutes post stress in the late phase). For average heart rate (BPM), there was a main effect of time (β=-0.11, SE=0.04, t-stat=-2.90, p=0.004), a stress by scan order interaction (β=-2.51, SE=0.81, t-stat=-3.08, p=0.002), and a time by scan order interaction (β=-0.12, SE=0.04, t-stat=-3.20, p=0.001). We also found a three-way interaction between stress, time, and scan order (β=0.20, SE=0.05, t-stat=3.83, p<0.001). Overall, participants who had the stress session first showed increased heart rate during the stress session, while we found no between-session differences in those who had the control scan first. Additionally, over time heart rate in the stress session remained higher than in the control session only in those who had the stress scan first (full model results in SM text 1).

There was also a significant main effect of time on heart rate variability (RMSSD, β=-0.13E-3, SE=0.052E- 3, t-stat=-2.494, p=0.013), a stress by time interaction (β=0.223E-3, SE=0.076e-03, t-stat=2.933, p=0.003), and a stress by scan order interaction (β=1.894E-3, SE=0.798E-3, t-stat=-2.373, p=0.018).

Follow-up tests indicated that participants who had the stress scan first showed reduced heart rate variability in the stress session (β=-0.003, SE=0.001, t-stat=-3.022, p=0.003, Figure 3). No significant effects were seen in participants who had the control session first. Additionally, over time, there was a significant increase in heart rate variability in the control, but not the stress session (see SM Text 1 for full results). Together, these results indicate that our stress induction procedure was successful in inducing the expected autonomic changes.

### Task-induced network activations in SN, ECN, and DMN

Before testing our main hypothesis with regards to temporal shifts in network activation under stress, we first verified whether the intermixed oddball, 2-back, and associative memory retrieval task blocks indeed induced the expected activation in SN, ECN, and DMN, respectively. In a GLM context, activity induced by each task was modeled in separate regressors and convolved with a canonical hemodynamic response function. For SN, we contrasted oddball trials with non-target faces, for ECN, we calculated the contrast between two-back target trials and non-target trials, while for DMN, we contrasted remembered and forgotten trials. See Figure 4 for whole-brain results.

**Figure 4.**
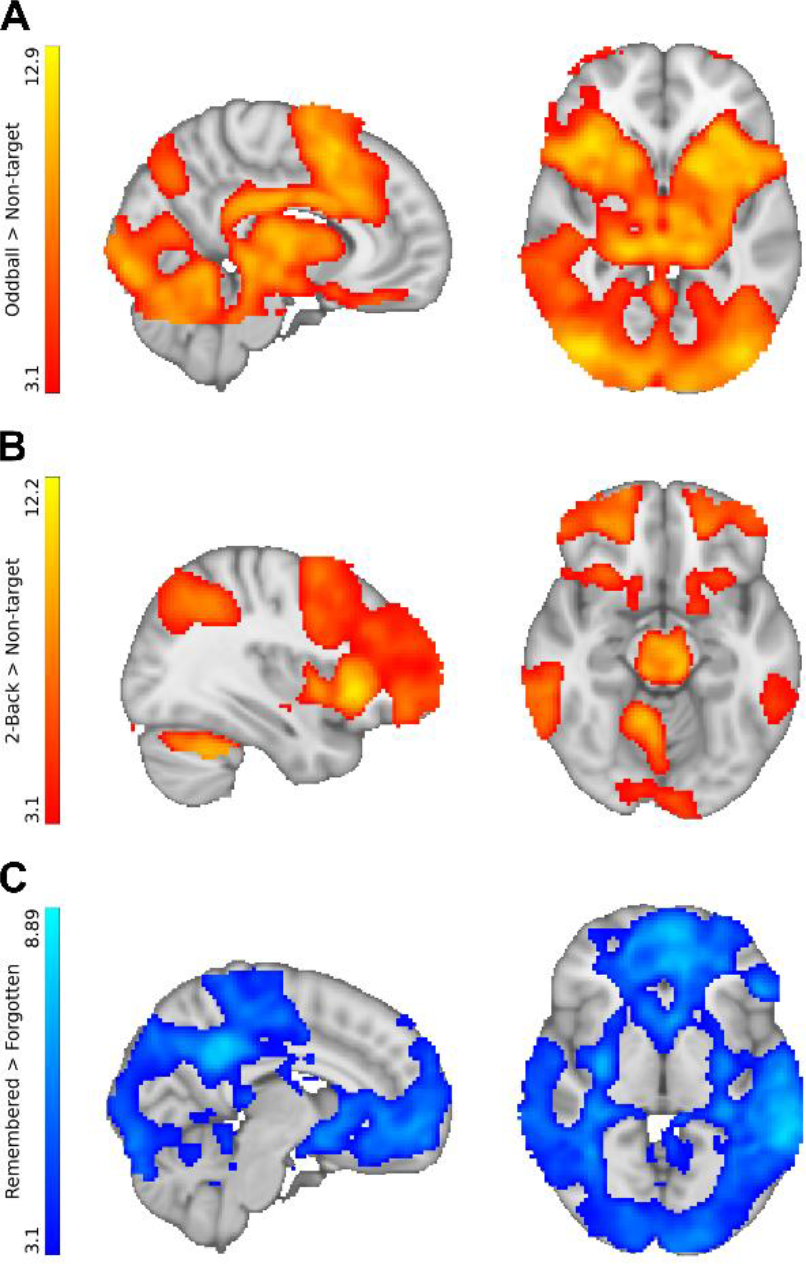
Main Task Effects. Main task effects across all task runs for each of the contrasts for the SN- oddball task (A, MNIxyz=41 91 35 mm), ECN-2back task (B, MNIxyz=27 33 29 mm), and DMN-associative retrieval task (C, MNIxyz=41 91 35 mm). Increased activity seen within expected regions of interest as well as some overlap in the SN and ECN contrasts, as well as an unexpected decrease in the DMN contrast. Results are whole-brain corrected with a cluster forming threshold of Z>3.1 and whole-brain corrected cluster significance level of P<0.05.

Contrast parameter estimates for each task were then averaged within each of the three networks and entered into mixed-effects models. As our hypothesis pertains to specific changes within networks over time, separate models were fit for the SN, ECN, and DMN. There was expected significant SN-related activity in the Oddball>Standard contrast (β=1.11, SE=0.06, t-stat=18.21, p<0.001) and significant activation in the ECN to the 2Back>non-target trials (β=0.57, SE=0.07, t-stat=8.73, p<0.001). Contrary to expectations, we saw a significant overall decrease in the DMN for the Remembered>Forgotten contrast (β=-0.35, SE=0.05, t-stat=-7.45, p<0.001).

### Stress results in shifts in SN and DMN

We next tested our main hypotheses regarding temporal shifts under stress in the networks across time in each network separately. Within the SN, there was a significant stress by phase interaction effect (β=0.22, SE=0.07, t-stat=02.99, p=0.003), with increased SN activity in the early phase of the stress response relative to the control condition (β=0.35, SE=0.13, t-stat=2.77, p=0.007), but not in the late phase (β=-0.10, SE=0.13, t-stat=0.77, p=0.444). Additionally, there was a significant reduction of activity in the late stress session compared to the early stress session (β=-0.44, SE=0.15, t-stat=-2.99, p=0.003). Within the DMN, there was an overall decrease in activity in the stress session compared to the control session across both phases (β=0.11, SE=0.04, t-stat=2.40, p=0.016). Furthermore, there was a significant main effect of scan order on DMN, with greater suppression seen in participants who had the stress scan first (β=-0.11, SE=0.05, t-stat=-2.24, p=0.025). No significant stress effects were seen in ECN activity (Main effect of stress: β=-0.02, SE=0.09, t-stat=-0.18, p=0.858, Figure 5A). We finally tested whether cortisol stress reactivity, as measured by the AUCi differences between stress and control sessions, moderated the effects of stress on any of the networks. No significant relationship between cortisol and any of the networks was found (analysis results can be found in the SM).

**Figure 5.**
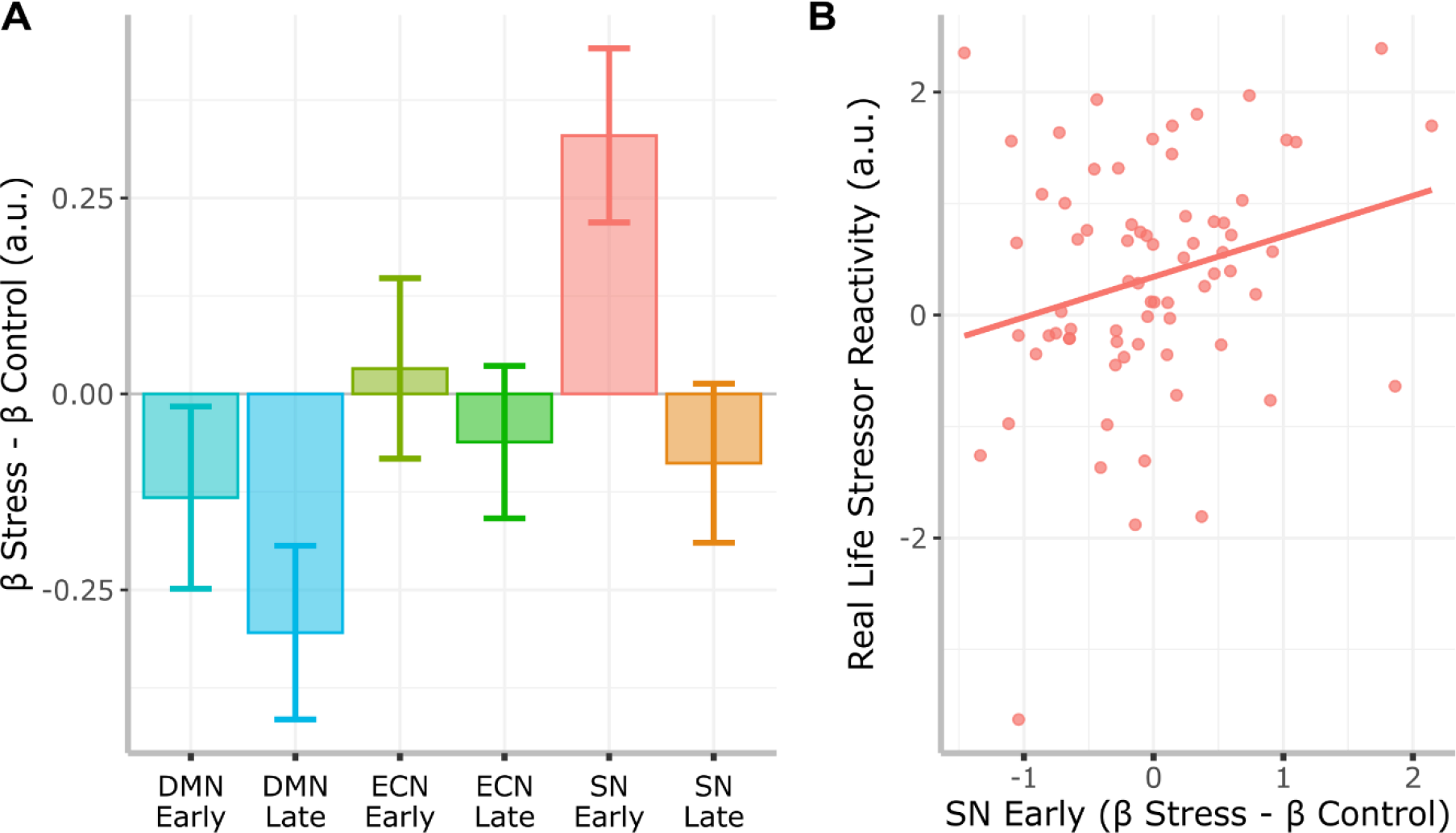
Effects of stress on large scale networks over time and real-life stress reactivity. A) Stress-Control differences in each of the networks in both the early and late phases of the stress response showing overall suppression of DMN under stress, and an increase in SN activity in early phase of stress reactivity. B) Decreased SN reactivity to stress is related to increased resilience in daily life. Error bars = SEM.

### SN stress reactivity is related to affective reactivity to stress in real life

We finally investigated the association between shifts in large-scale networks under stress and real-life affective reactivity to stress using mixed effects models with network as a factor. There was a significant three-way interaction between stress, phase, and real-life affective reactivity within the SN, but not the ECN and DMN, indicating that affective reactivity in real life is linked to timing-dependent changes in the central response to stress (β=-0.23, SE=0.10, t-stat=-2.17, p=0.030). Follow-up tests showed that effects were driven by changes in the early phase of the stress response (β=0.71, SE=0.35, t-stat=-2.05, p=0.045) as opposed to the late phase (β=0.18, SE=0.35, t-stat=0.50, p=0.622; Figure 5B).

### Stress related SN activity enhances vigilance

We finally investigated associations between changes in networks under stress and performance measures from each of the corresponding tasks. This was done to investigate the impact of stress not just on brain activity, but also on task performance associated with the networks. In the oddball task, participants reacted significantly faster to the presentation of an oddball stimulus in the stress compared to the control session (β=0.97, SE=0.01, t-stat=-3.09, p=0.002). There was an additional main effect of time, with average reaction times being slower in the late phase of the MRI scan (β=0.98, SE=0.01, t-stat=-4.09, p<0.001). An exploratory mediation analysis was run to examine whether this was due to significant stress effects in SN under stress. Mediation models were run with simple slopes and subject as a random effect. Stress-induced SN activity significantly mediated the relationship between stress and reaction times during the presentation of oddball stimuli (β=0.0361, 95% CI=[-0.819, 0.00], p=0.03). This indicates that increased SN activity under stress enhances reaction times in a vigilance-oriented task.

Participants furthermore performed worse on the oddball facial recognition task outside of the scanner (immediately after the early phase scans, and then again the later phase) in the stress versus the control session, as measured by d-prime (β=-0.42, SE=0.06, t-stat=7.55, p<0.001, Figure 6A). This was driven by both fewer hits (β=-3.29, SE=0.67, t-stat=4.87, p<0.001, Figure 6B), and more false alarms (β=1.68, SE=0.42, t-stat=4.04, p<0.001, Figure 6C). Interestingly, there was also a significant interaction effect of session (stress versus control) and SN activity during encoding on subsequent false alarms (β=1.05, SE=0.48, t-stat=2.18, p=0.030, Figure 6D). Post-hoc tests contrasting stress and control sessions showed a significant difference between slopes (β=-1.05, SE=0.5 t-stat=-2.098, p=0.0371), with a steeper positive slope of the regression of false alarm rates onto SN activity for the stress (slope=0.461) versus the control condition (slope=-0.588).

**Figure 6.**
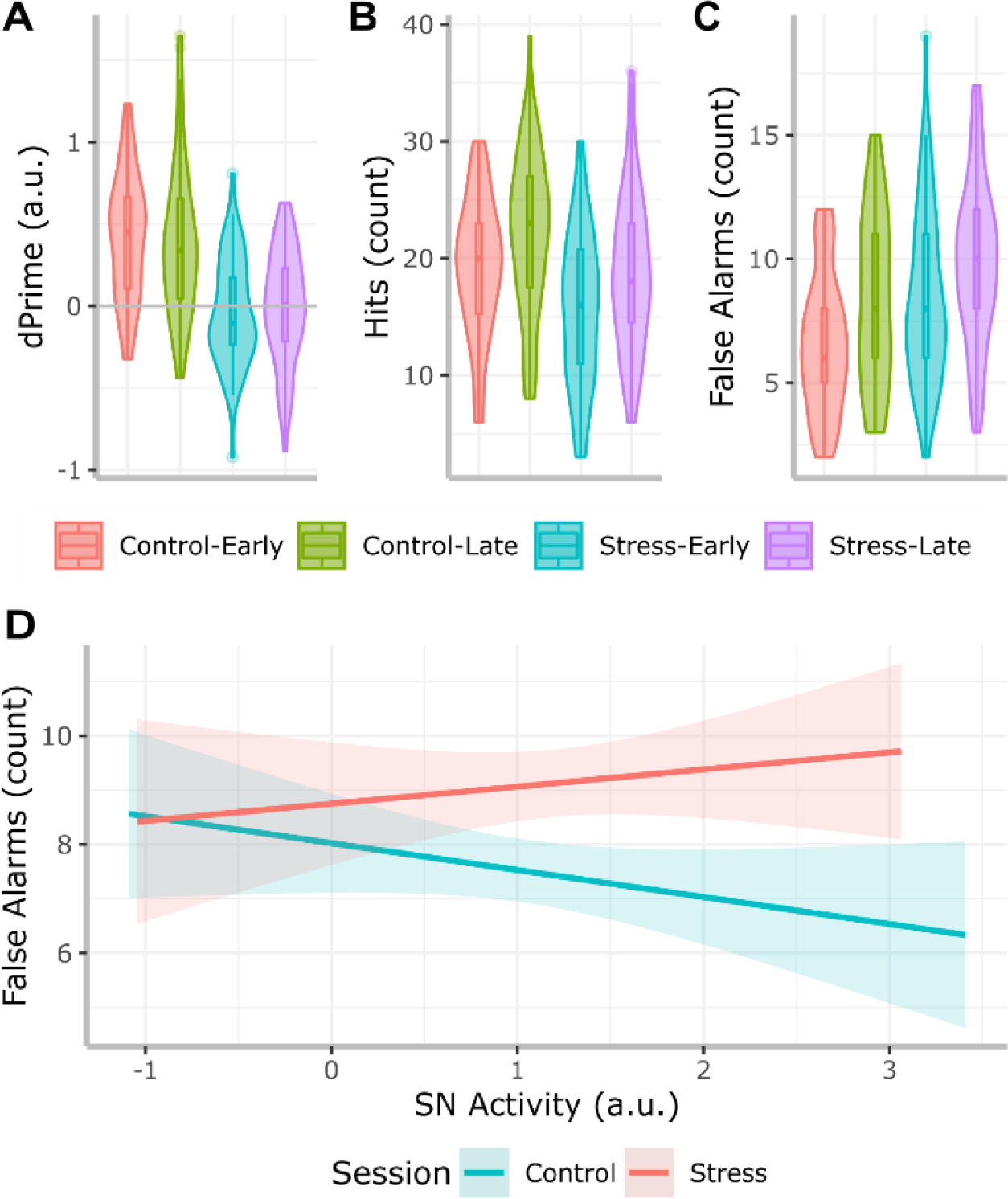
Oddball Task Measures. D-prime (A), Hits (B), and False alarms (C) differed significantly between the stress and control sessions, with lower d-prime under stress driven by both fewer hits, and more false alarms. SN (salience network) activity was also related to false alarms differently in the stress and control session (D).

A one sample, two-sided t-test against the expected chance level (25%) in the DMN associative memory retrieval task (performed during scanning) showed that participants performed significantly above chance levels (Mean accuracy=61.5%, t-stat=255.65, p<0.001). There were significant stress effects or phase effects neither in reaction times, nor in the ability to recall images. There was, however, a significant effect of valence. Participants responded faster (β=-0.0018, SE=0.0009, t-stat=-2.063, p=0.0392) and with more accuracy (β=0.027, SE=0.004, t-stat=7.266, p<0.001) to images that were negatively valenced compared to neutral ones. There were no significant differences between stress and control sessions in the 2Back/ECN task when looking at the reaction times, proportion of errors, or the LIASES score (Null results in online notebook).

## Discussion

In this study, we aimed to test how brain activity changes over the course of the stress response, and whether changes in large-scale neural networks under stress are associated with real-life stress reactivity. Laboratory stress induced a network-dependent and time-dependent shift in BOLD activity, with an early increase and a later decrease in responsiveness of the salience network (SN), alongside increased suppression of the default mode network (DMN) throughout. Importantly, increased early SN reactivity to stress was associated with increased affective reactivity to stress in real life as measured with ecological momentary assessment (EMA). Our results highlight how neural network configurations time-dependently change in response to (laboratory-based) stress induction. Crucially, interindividual differences in these network reconfigurations are related to affective reactivity to real-life stressors.

Affective reactivity to daily-life stress was linked to SN reactivity to salient stimuli in the early, catecholaminergically dominated, phase of the stress response. Although we found a return to baseline, we found no association with SN activity during the late phase of the stress response, which is thought to be dominated by genomically driven effects of glucocorticoids. Interestingly, our affective reactivity measure corresponds to in-the-moment stress, which falls into a time scale similar to that of the early phase of the acute stress response. Our results are in line with two previous studies that connected real-life affective dynamics to neural measures. One study was able to link negative affect inertia to increased responses to social feedback in the insula – a core node of the SN ^32^. Another study found a link between stress-induced dopaminergic activity using PET to psychotic reactivity to stress in real life ^33^. The usage of PET imaging however does not allow for investigations of temporal shifts as a result of hormonal changes relating to stress response. Utilizing fMRI, we were able to link real-life measures to the full scope of the stress response, including both the reactivity and recovery phases.

In our data, stress-related SN activation mediated decreased reaction times to unexpected oddball stimuli in the stress session, indicating heightened vigilance supported by SN activity. Previous evidence has linked the SN to vigilance and attentional reorienting mechanisms^34,35^ which are enhanced by stress^36^. As suggested by our findings, in real-life, this may translate to increased vigilance in stressful situations, resulting in greater affective reactivity to stressors. This mechanism may possibly be related to increased attention to threats or negative events, and may be at the core of attentional bias to negative or stressful events typically observed in anxiety and depression^37^.

The fMRI results additionally confirm the hypothesis based on the model proposed by Hermans and colleagues (2014) that the early stress response is characterized by a shift towards increased SN activity in a task-based setting^1^. This is in line with findings of increased functional connectivity in the SN in response to stress^7,38,39^, a process that is thought to be driven by norepinephrine release from the locus coeruleus (LC)^3,12^. Norepinephrine release is accompanied by activation of the sympathetic autonomic nervous system, resulting in increased heart rate, which often coincides with reduced heart rate variability^15^, as also found in our study. Finally, we found a return to baseline in SN activity in the later recovery stage of the stress response where corticosteroids have been shown to be involved^1,17^. This indicates that the initial shift in stress systems is later reversed when recovering from a stressor.

We found suppressed DMN activity during the early phase of the acute stress response, which is in line with previous studies^16,40^. More surprisingly, this suppression persisted in the recovery phase, two hours after the onset of the stressor. The DMN is implicated in memory retrieval functions, as well as in self-referential processing and rumination^41–44^. Yet, no impairment of retrieval functions as a function of stress was seen on the associative memory retrieval task conducted during scanning. We did however observe a negative effect of stress on the oddball facial recognition performance just after scanning.

However, this effect is difficult to interpret because encoding for this task also took place after stress induction. Worth noting is that the reduction in DMN may instead indicate decreased internally directed cognition, which is the cost of increased SN-driven exogenous attention in the early phase^42,45^.

Additionally, no links between the DMN and real-life stress were established. While previous evidence has linked the DMN to depressive rumination and negative affect in daily life^44^, our findings do not support the role of DMN-driven rumination as a mechanism of affective reactivity to stress.

It is worth noting that the associative retrieval task did not elicit increased DMN activity relative to a fixation baseline, in contrast with previous evidence^41,45–47^. Our specific paradigm may not allow for sufficient DMN engagement given the traditional view of the DMN as a task negative network^48^. Another possible explanation may come from a deviation from the original protocol which used only neutral images^49^. Our design utilized both negative and neutral images, which is a commonly used procedure to investigate affective processes^50,51^. Given the higher accuracy and faster reaction times to negatively valent images, it may be that valence-related mechanisms overshadowed previously reported task effects, resulting in suppression of the DMN in our contrast. Despite that, however, we were still able to see stress effects on DMN in our study.

We expected to see decreased ECN activity in the early reactivity phase of the stress response, and an increase in the late phase ^1,15^. This latter process is putatively driven by genomic effects of cortisol released in the early phase ^4^. Previous work has shown coupling of ECN and SN under stress, with a breakdown of this process only at higher levels of arousal ^39,52^. It is possible that arousal levels in our study were not high enough to induce this decoupling, thus resulting in co-activation of ECN along with SN. This weaker stress response may also explain the lack of ECN upregulation during the late phase, as it is thought to occur due to genomic influences of cortisol released in the early phase ^4^. Previous work has found administration of corticosteroids enhances ECN activity within the time frame of genomic corticosteroid effects following stress exposure^19,53^. Such studies used high doses of corticosteroids (peak measures of salivary cortisol at around 40-130 nmol/L) that may not be comparable to those resulting from the stress induction paradigm (around 5.6 nmol/L in our study). Higher doses in pharmacological studies, as well as individual variations in responses to stress induction may explain the lack of suppression. Finally, we found a significant sex difference in cortisol responses that was driven by blunted responses in males relative to hormonal contraceptive users. While important to investigate, such effects of contraceptive use and sex are beyond the scope and power of the current study, and thus we refrain from further interpretation.

In addition to neural findings, we also demonstrate the successful utilization of a dynamic EMA-based residualization method to a daily affect measure. This measure is novel in capturing interindividual differences in positive affect changes in response to in-the-moment stress. We focus on positive affect as our previous work has shown that stress seems to have a bigger impact on positive rather than negative affect^54^. This may be due to respondents showing greater variability when answering positive affect items, which is reflected in the skewed distribution of negative affect. Additionally, previous work has linked self-reported momentary resilience directly to positive affect in daily life^30^. By correlating this measure to neuroticism, we also show that it partially relates to an established personality trait that is linked to psychopathology^55^.

While our results on stress dynamics in the lab and real life are novel, their long-term relationship to resilience is still an open question. Some studies have shown affective responses in daily life to be linked not only to momentary mood and psychiatric symptom expression, but also future mental health outcomes^56–58^. Thus, our measure may capture these same mechanisms at work in a shorter time frame. Some prospective studies have also investigated how long-term resilience is related to brain activity of large-scale networks. These studies have shown increased SN activation to be predictive of later mental health outcomes as well^38,59^. Together, these findings suggest that the mechanism leading to poorer mental health outcomes could be related to stress-induced enhancement of vigilance related processing that has a long-term impact.

In conclusion, our study demonstrates that neural stress reactivity is associated with stress reactivity in real life. These findings demonstrate ecological validity of neuroscientific research in the context of stress. Furthermore, partly in line with our hypothesis, we demonstrate a change in large scale networks under stress, with increased SN reactivity immediately following threat that returns to baseline during stress recovery. Importantly, our results indicate that SN-related attentional mechanisms following stress are linked to affective reactivity in daily life. Individuals who show enhanced SN activity immediately following a laboratory stressor also show enhanced reactivity to stressful events in real-life contexts outside of the laboratory. Mechanisms such as increased vigilance under stress may have implications for our understanding of how stress related disorders develop. It may be that increased vigilance to threat can lead to impaired recovery from stress in the long-term, resulting in poorer mental health outcomes. Indeed, recent studies have indicated this hypervigilance may be risk markers for development of stress related disorders ^60^. Future extension of these findings in prospective studies and clinical populations may help in uncovering the real-life consequences of neural underpinnings of these disorders.

## Methods

### Participants

Eighty-three students in the first year of the bachelor’s program for biomedical science and medicine were recruited for this study. Participants were healthy, right-handed, Dutch speaking volunteers with no history of psychiatric or neurological illness at the time of recruitment. Participants first completed two counterbalanced weeks of stress assessments in daily life (i.e., Ecological Momentary Assessments, EMA), one with an ecological exam stressor and the other without. At the end of each of the weeks, participants completed a series of computer tasks and a questionnaire battery. This was followed by two counterbalanced fMRI sessions, one with a stress induction procedure using a modified version of the socially evaluated cold pressor test (SECPT, Schwabe & Schächinger, 2018), and the other with a matched control task. All procedures were approved by the local medical-ethical committee (METC Oost-Nederland; Figure 1).

Five participants withdrew prior to completion of all scanning sessions. Due to issues that occurred during scanning, an additional three participants were also excluded. Reasons included incorrect stimulus presentation during scanning (1), scanner-related malfunctions (1), and incidental findings (1). Finally, due to the COVID-19 outbreak an additional five participants were unable to complete either one or both scan sessions and were thus excluded from the MRI portion of the study, bringing the total number of participants to 70 (f=43 (61.4%)).

### Real-life affective reactivity

Participants completed repeated EMA surveys (or beeps) delivered to their phones six times a day for two separate weeks: One week during a high-stakes examination period (i.e., a stress week), and the other outside this period (i.e., control week). Surveys assessed stress levels as predictors and affect as an outcome. Stress was assessed using questions regarding (1) the most stressful event experienced since the last beep (i.e., event-related stress), (2) stress related to the activity participants were engaged in when answering the surveys (i.e., activity-related stress), (3) stress relating to the social context participants were in at the time of the beep (i.e., social stress), and (4) physical stressors. Affect items were collected for positive and negative mood. Additionally, ambulatory data was collected from wrist-worn devices measuring aspects of physiological arousal. At the end of each of these weeks, participants filled in a questionnaire battery. Within the scope of the current paper, only subjective stress and positive affect measures are used from the EMA data. Full details and results of the EMA weeks are reported in previous work, and the full questionnaire set can be found in the associated GitHub directory^54^.

### Laboratory stress and the neural response

We investigated laboratory stress using a multi-day fMRI paradigm. Participants took part in three MRI scan days: one structural scan day and two functional scan days (stress and control days, order counterbalanced between participants). Structural scans were used to reduce scanner-related apprehension in scanner-naïve participants. Those with prior scanning experience were only scheduled for the two functional scan days, and structural scans were appended to the end of their first fMRI session. Participants were asked to be present two hours prior to the functional scans, between 10:00 and 16:00, to allow cortisol levels to return to baseline prior to testing, and to account for diurnal cortisol fluctuations. During this time, participants practiced the tasks they would later perform in the scanner (reported in the fMRI Task Section, Figure 1C). Following the rest period, participants were escorted to the MRI scanner, where skin conductance electrodes and a PPG heart rate sensor were attached to their left hand, and respiration belt was attached below the chest. The stress sessions included a modified version of the Socially Evaluated Cold Pressor Test (SECPT), and the other a matched control protocol ^61^.

The SECPT or control procedure began promptly following scanner set-up. The SECPT was performed by a novel male experimenter with whom participants had no prior interaction (i.e., the stressor). The stressor briefly introduced himself and proceeded with placing the participant’s foot in ice water (0-2 °C) for three minutes. Participants were able to ask to have their foot removed earlier, but only if they found the procedure unbearable. This was followed by a mental arithmetic task where participants were required to count backwards from 1872 (the number of contractual hours the authors work per year) in steps of 17 for three minutes. If they were too slow, or made a mistake, the stressor would ask that participants restart the task. The stressor maintained a neutral tone, and eye contact with the participant throughout. The control session was matched for time, with a familiar experimenter performing all procedures. For this procedure, water was at room temperature (21-25 °C), and the arithmetic task was to count upwards from 0 in steps of 5. Participants were not corrected in the event of errors. At the end of the stress session, participants were debriefed regarding the SECPT protocol.

Saliva samples were acquired before and after the stress/control procedures. A full SOP is provided in the data repository.

Saliva sampling began before stress induction (T1=-3min). This was followed with a saliva sample after stress induction (T2= 14 minutes). The remaining samples were acquired immediately following the first resting state run (T3=42 minutes), at the end of the early phase of scanning (T4=87 minutes), and at the end of the last phase of scanning (T=160 minutes). Saliva samples were stored on site at -80° C until they were shipped for offsite analysis in Dresden, Germany. After thawing, salivettes were centrifuged at 3,000 rpm for 5 min, which resulted in a clear supernatant of low viscosity. Salivary concentrations were measured using commercially available chemiluminescence immunoassay with high sensitivity (IBL International, Hamburg, Germany).

### fMRI Tasks

Participants performed a series of fMRI tasks in two phases immediately following the SECPT/Control procedure. The first phase examining stress reactivity (i.e., the early phase) followed the SECPT/control procedure and consisted of three task runs of 11.5 minutes each, with two interleaved resting-state runs of five minutes in between (Figure 1B) for a total of approximately 50 minutes including saliva sampling following the first resting-state run. Participants were then taken out of the scanner for a short 20- minute break before continuing with the second half of scanning (hereafter the late phase). A total of 21 different task blocks were presented during each run, and a total of 21 blocks of each task were shown in each phase of the scanning session. Task blocks consisted of 27 seconds, followed by 6 seconds of rest before the next task. During this time, participants were shown a shape on the screen that indicated what the next task would be. Task order was pseudorandomized to maximize the number of times participants had to switch from one task to another. Tasks were selected based on previous evidence implicating the involvement of one of the three networks of interest (i.e., SN, ECN, DMN) as follows:

### Oddball

A facial oddball paradigm was selected for SN recruitment based on previous research showing robust SN activity during the presentation of an oddball stimulus^35^. Participants were shown a stream of faces with one standard neutral face being shown most of the time, and a novel oddball face with an emotional expression being shown 16% of the time. Participants also performed a facial recognition task outside the scanner following each phase (i.e., early and late), with 40 oddball targets shown in the scanner, and 20 lures. Faces were selected from four databases: Amsterdam Face Database, Radboud Face Database, Karolinska Directed Emotional Faces, and the Chicago face database ^62–65^.

### 2-Back

Working memory tasks are often used to recruit the Executive Control Network ^66^. To this end, we selected a 2-back task. Participants were presented with a stream of numbers and were instructed to press a button when the most recent number they had seen was the same as the number two places back (e.g., 3, 1, 3). A total of 15 trials per block were presented. Individual trials lasted for 1.8 seconds, with a stimulus being present for approximately 400 milliseconds ^15^.

### Associative Retrieval

The default mode network has been shown to be active during internal thought processes, and memory retrieval tasks. To this end, we used an adapted version of an associative retrieval task ^49^. On the scan day, but prior to the scan, participants performed two encoding sessions spaced 45 minutes apart to simulate the time in between the two scan phases (i.e., early and late encoding) during that day. During the encoding sessions, participants viewed a series of images that were negatively or positively valent which moved to one of four corners of the screen. Participants were told that they would be tested on the object-location association in the scanner. During scanning, participants engaged in a retrieval task where they were presented with the images from the encoding session and asked to indicate the location of the image using a button box.

### EMA Data Analysis

We adopted a residual-based stress score to derive a measure of real-life affective reactivity to stress ^25^. First, two general linear mixed effects models were used to estimate the effect of stress exposure (i.e., control or exam week) on stress reports (model 1) and positive affect (model 2). For model 1, an aggregate measure of stress was calculated from the total of the event, activity, and social stress scales. When participants reported stress levels above their individual means, surveys were labelled as “More Stress”. Responses below the individual mean were labeled as “Less Stress”. This was done to reduce affect-congruency effects and was modeled using a binomial family. Model 2 used positive affect as a continuous variable. In both models, we added a random effect of subject, and a random slope and intercept for week type effects. Random effects of these models would therefore indicate the subject level impact of stress on subjective positive affect. Random effects were then modeled against each other using linear regression to derive the overall change in positive affect relative to experienced stress during the stress week. Residuals from these models were used as a residual-based affective stress reactivity measure. In order to make interpretation easier, inverse scores are presented where increased scores represent increased affective reactivity in real life. This score was used in the fMRI models described below.

### fMRI Arousal Measures

We next established the validity of our laboratory stress induction procedure by examining the effects of stress induction via the SECPT on biophysiological measures of salivary cortisol and heart rate (IBI and RMSSD) using mixed effects models with subject as a random effect and session (i.e., stress or control), time, and scan order as fixed effects. Sex was controlled for as a fixed effect. Interactions were modeled for all three of these predictors. Session was modeled with a random slope and intercept. Distributions of the data were examined, and model families were chosen according to the optimal fit as determined by residual normality and the Akaike Information Criterion (AIC). Follow-up models were carried out using the “emmeans” package. The difference in the area under the curve with respect to increase (AUCi) for the cortisol response was calculated per subject using the first four samples to derive individual measures of cortisol stress reactivity (AUCistress – AUCicontrol).

### fMRI Preprocessing

Preprocessing was carried out for each run separately. DICOM images were first converted to NIFTII format. Echoes for each run were realigned to the first acquired volume and then recombined using PAID weighting method ^67^. 3D images were then temporally concatenated into a single 4D file. Additional processing was done using FSL’s FEAT (FMRI Expert Analysis Tool, version 6.0.0). Registration to high resolution structural and standard space images was carried out using FLIRT ^68,69^. Registration from high resolution structural to standard space was then further refined using FNIRT nonlinear registration.

Motion parameters were estimated using MCFLIRT, and interleaved slice-timing correction applied using Fourier-space time-series phase-shifting. Non-brain removal was carried out using BET and spatial smoothing using a Gaussian kernel of FWHM 6.0mm; grand-mean intensity normalization was carried out by a single multiplicative factor. Motion artifact removal was then carried out using non-aggressive AROMA, followed by high pass filtering of the data ^68,70,71^.

Task regressors were modelled for each event in our three tasks. In order to keep contrasts clearer, only contrasts within the same task were used to model task-related activity. For the 2-Back task, a regressor was modeled for the occurrence of 2-Back target events, and for the non-target events (i.e., No-Backs) with a 0.3 second duration. For the oddball task, oddball and non-oddball events were modeled as separate regressors with 0.3 seconds durations each. For the memory task, the later remembered and forgotten trials were each modelled as separate regressors, with durations set to 3.5 seconds. Specific contrasts based on these regressors were used to model activity within three networks of interest: The Executive Control (ECN), Default Mode (DMN), and Salience (SN) networks. To model ECN activity, the 2- Back events were contrasted against the No-Back events (i.e., ECN=2-Back>No-Back). Salience network activity was modelled by modelling the oddball events against the non-oddball events (i.e., SN=Oddball>Non-oddball). Finally, to model the DMN network, the contrast for both remembered vs forgotten images used. Time-series statistical analysis was carried out on the contrasts of interest using FILM with local autocorrelation correction. Z (Gaussianised T/F) statistic images were thresholded non-parametrically using clusters determined by Z>3.1 and a (corrected) cluster significance threshold of P=0.05 ^72,73^. Results of the tasks from each run were analyzed at the subject level using a fixed effect model in FEAT.

### fMRI Data Analysis

First level fixed effects models were then constructed at the run level. For the 2-Back task, we modelled the occurrence of a 2-Back target and the non-targets as events and the contrast 2-Back>non-target used to model ECN activity. For the oddball task, oddball and standard trials were each modelled as events.

The contrast used to measure SN activity was the Oddball>Standard trials contrast. Finally, for the associative retrieval task, remembered and forgotten trials were each modeled, and the contrast Remembered>Forgotten was used to model DMN activity.

Second-level analysis was performed using a 2x2 fixed effect design, with session (control vs stress) and phase (early: runs 1-3 and late: runs 4-6) as fixed effects resulting in four main EVs: Control Early, Control Late, Stress Early, Stress Late. Contrasts were modelled for mean task-related activity across the four EVs. Z-statistic images were thresholded using non-parametric cluster-based thresholding at Z>3.1 and a corrected cluster significance threshold of p=0.05. Previously established templates were used for ROI analyses, with a single mask created from all subdivisions of the SN, ECN, and DMN ^74^. Threshold z-statistics were extracted from specific target task-network combinations for each of the four levels (Control Early, Control Late, Stress Early, Stress Late).

We first investigated the main effects with three-way interactions for session (stress/control), phase (early/late), and network (SN/ECN/DMN). We next investigate the specific hypothesis regarding temporal shifts as a function of stress in each network individually, with additional interaction terms modeled for cortisol stress reactivity and the real-life affective reactivity score. A maximal fitting approach was used in which subject was modeled as a random effect, and correlated random intercepts and slopes modeled for all other fixed effects of interest. Since AUCi and the real-life stress score only have one level per subject, no random slopes or intercepts were added for these terms. Scan order was also modeled as a fixed effect with no random slope and intercept to control for potential differences due to the order of sessions.

### Behavioral Task Analysis

Task-related data was analyzed using mixed-effects models, with individual trials being modelled for each of the three tasks. Trials with reaction times lower than 200ms were removed prior to analysis. For the 2-back, reaction time, error count, and a combined score (LIASES score) were analyzed^75^. For the oddball task, the average reaction time per trial and the oddball facial recognition (as measured by dprime, *d’*) and the number of hits and misses were analyzed. For the associative retrieval task, reaction times and recall percentages were analyzed. In all models, session (i.e., stress or control MRI) were modelled as fixed effects with an interaction term for phase (i.e., early or late). A random effect was modelled for subject, with a correlated random intercept and slope modelled for session and phase effects. The same models used for the reaction time were used for the out-of-scanner oddball recognition task. We additionally ran separate models with an interaction term modelled for neural activity in the targeted ROI’s to examine the relationship between task performance and neural responses, with random slopes and intercepts also modelled for ROI activity.

## Online Notebook and Data Availability

Data is made available via our institutional repository upon request due to ongoing usage of the data. Due to the usage of multi-echo imaging and the associated large storage requirements, only recombined Niftii files are made available, and not raw DICOMS. Skull stripped T1 images are made available for anonymization reasons. All scripts are publicly available on GitHub (https://github.com/raytut/Large-Scale-Networks-and-Real-Life-Stress), along with an analysis notebook containing the results produced in R (Version 3.6.1).

## Acknowledgements

We would like to thank Tim Vriens and Bob Kapteijns for their assistance in data collections. This work was supported by the European Research Council (ERC2015-CoG 682591).

## Author Contributions

Conceptualization: RT, EH, MK, FK

Methodology: RT, EH, MK, FK

Data Collection: RT, NK

Data analysis: RT, EV, EH

Writing: RT, EV, EH

Editing: All authors Funding: EH

