## Supplementary material for "Changes in large-scale neural networks under stress are linked to affective reactivity to stress in real life": SM

| **SM Table 1. Individual task details** | | | | | |
| --- | --- | --- | --- | --- | --- |
|  | ***Block Duration(s)*** | ***Trials per block (n)*** | ***Targets per block (n)*** | ***Trial Length (ms)*** | ***ISI  (ms)*** |
| ***SN: Oddball*** | 27 | 12 | 1-3 | 2200 | 1350-1450 |
| ***ECN: 2-Back*** | 27 | 15 | 2-4 | 1800 | 1350-1450 |
| ***DMN: Associative Retrieval*** | 27 | 3 | 3 | 9000 | 3500-7000 |
